# Variation in season length and development time is sufficient to drive the emergence and coexistence of social and solitary behavioral strategies

**DOI:** 10.1101/2024.06.18.599518

**Authors:** Dee M. Ruttenberg, Simon A. Levin, Ned S. Wingreen, Sarah D. Kocher

## Abstract

Season length and its associated variables can influence the expression of social behaviors, including the occurrence of eusociality in insects. Eusociality can vary widely across environmental gradients, both within and between different species. Numerous theoretical models have been developed to examine the life history traits that underlie the emergence and maintenance of eusociality, yet the impact of seasonality on this process is largely uncharacterized. Here, we present a theoretical model that incorporates season length and offspring development time into a single, individual-focused model to examine how these factors can shape the costs and benefits of social living. We find that longer season lengths and faster brood development times are sufficient to favor the emergence and maintenance of a social strategy, while shorter seasons favor a solitary one. We also identify a range of season lengths where social and solitary strategies can coexist. Moreover, our theoretical predictions are well-matched to the natural history and behavior of two flexibly-eusocial bee species, suggesting our model can make realistic predictions about the evolution of different social strategies. Broadly, this work reveals the crucial role that environmental conditions can have in shaping social behavior and its evolution and underscores the need for further models that explicitly incorporate such variation to study evolutionary trajectories of eusociality.

## Introduction

Environmental conditions can have a major impact on the expression of social behaviors. Variation in the social structure of animal groups has been documented across both latitudinal and altitudinal gradients in birds (1), bees (2-5), ants (6-7), wasps (8), and even social spiders (9-10). This variation is tightly linked to changes in season length and associated variables such as temperature and resource availability; these interrelated variables can alter the costs and benefits associated with social living (1,6,9,11). For example, changes in temperature can influence how quickly offspring develop (12), when individuals can forage (13), and the availability of food resources in the surrounding environment (14-15). In turn, changes in seasonality have dramatic effects on the behavioral strategies favored by individuals – for instance, paper wasps with a high level of food availability are more likely to delay reproduction (16).

Eusociality represents one of the most extreme forms of social living whereby reproductive individuals live and cooperate with nonreproductive workers to reproduce as a group (17-19). The transition from individual to eusocial reproduction is often considered one of life’s major evolutionary transitions (20), and eusocial groups have arisen multiple times throughout the animal kingdom, including multiple origins in both insects and vertebrates (21). Some of the best known examples of eusociality are found among the social insects (Insecta:Hymenoptera), which include eusocial bees, ants, and wasps.

There is a rich history of models that study the set of preadaptations necessary before eusociality becomes an evolutionarily stable strategy (16-17, 22-23). These models have helped to reveal the life history traits and ecological conditions which favor the evolution of eusociality and help to explain the multiple independent origins of eusociality across different taxa. For example, work by Seger (17) and Quiñones and Pen (23) have demonstrated the importance of considering the temporal structures of life histories in the emergence of eusociality. Seger (17) modeled life history as a process where reproductive females can produce multiple, overlapping broods per season (e.g. partially bivoltine). Through the incorporation of these multiple broods, the model reproduced the sex ratio biases observed in natural populations of bees and wasps. Quiñones and Pen (23) later expanded on Seger’s framework to identify sets of preadaptations that can trigger the transition from a partially bivoltine, solitary life history to a eusocial one, including: haplodiploidy, maternal control over sex ratios, adult diapause (e.g. overwintering, mated females), and the presence of a protected nest site. Together, these models have provided a framework for understanding which life history transitions are associated with the evolutionary origins of eusociality in social insects. However, both models treat each season as two generations during which the reproductive female’s behavior is invariant, limiting the ability to make inferences about the costs and benefits of social behavior under different environmental conditions.

Here, we extend these models by incorporating two additional conditions that can significantly impact the emergence and maintenance of different behavioral strategies: variable season length and differing offspring development times. Through the integration of these two parameters, our individual-centered model thus enables a direct examination of how variation in seasonality can shape the costs and benefits of social living.

Our model is based on the life histories of halictine (sweat) bees, though it could easily be adapted to a wide range of life histories. Halictines encompass a wide range of social behaviors, from solitary to eusocial (24-26). Throughout their evolutionary history, eusociality was gained at least twice (25) and there were also many subsequent reversions back to a solitary life history (24). As a result, this group of bees varies naturally and extensively in their social structure both within and between species (24-27).

Solitary halictines typically produce a single brood of offspring that contains a mix of reproductive males and females (26), though many solitary halictines do produce multiple broods of reproductives throughout the course of a breeding season. Solitary offspring that emerge early in the season can often produce their own reproductive generation later in the season, leading to multiple solitary generations within the course of a single breeding season. In contrast, eusocial sweat bees first produce a nonreproductive, worker brood followed by a larger brood of reproductives. In all temperate eusocial females, reproductive females mate and overwinter as adults before founding new nests the following season. Phylogenetic studies indicate that the evolution of adult diapause precedes the origins of eusociality in bees (28); thus, each season in our model is initiated by overwintered, mated adult females.

We modeled the ratios of male to females and helpers to reproductives as alleles which can change both day-by-day (during the development of the colony) and year-by-year (over evolutionary time), thus allowing us to examine a full range of solitary and eusocial behavioral strategies. We then used the evolving variation within these alleles to investigate how season length impacts the success of social and solitary reproductive strategies. Using these results, we predict the environmental conditions where eusocial and solitary strategies are expected to occur in nature, and we compare these predictions to species occurrence data for a group of socially variable bees, thus allowing us to link our theoretical model to naturally occurring variation in eusociality.

## Materials and Methods

To gain insight into the effects of season length on solitary-social transitions, we developed a stochastic, individual-focused approach adapted from Quiñones and Pen’s model of hymenopteran preadaptations (23). Our model expands on this and Seger’s (17) but differs by allowing both sex ratio and voltinism to emerge freely as well as by treating the foraging season as a series of discrete days, rather than bouts of reproduction. These modifications allow us to directly capture the relationship between season length and the emergence of eusociality. We ran this model until trait values stopped changing substantially over time, and we observed the emergence of both solitary and eusocial reproductive strategies that share many similarities with described nesting strategies in bivoltine insects, including halictine bees (Figure 1a; 26-27).

**Fig. 1.**
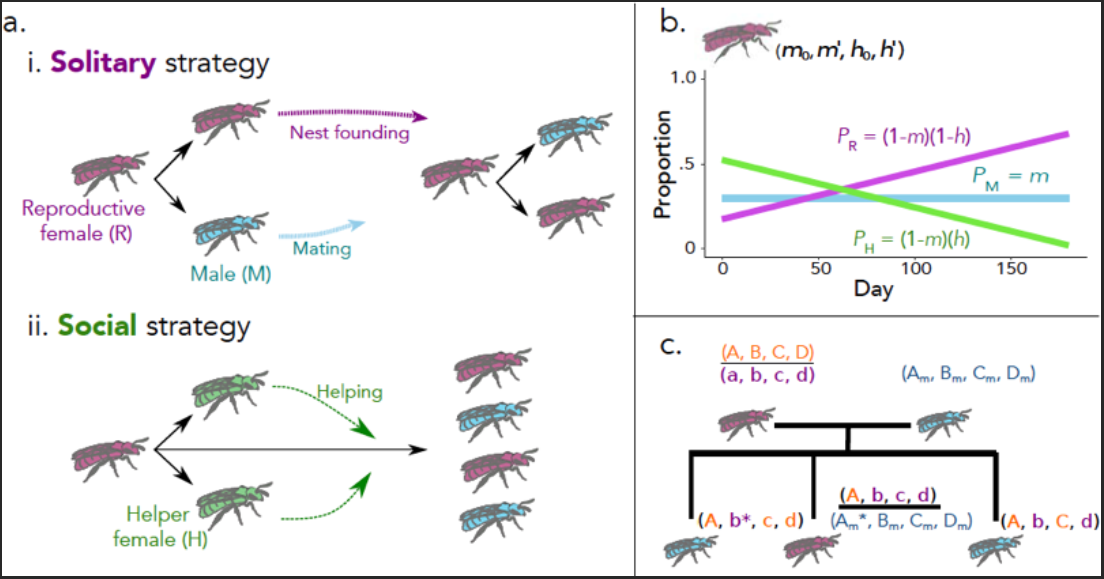
Illustration of the theoretical model. (a) Schematic of the two primary behavioral strategies arising from this model. (i) Solitary reproductives primarily produce reproductive males and females, at a total rate of *α* bees per day. Mature reproductive females (“queens”, purple) eventually form their own nests, producing their own offspring. Reproductive males (blue) leave the nest as soon as they mature, fertilizing females from all nests. (ii) Social individuals additionally produce female helpers (green) which never leave the nest, nor have offspring of their own. Rather, through helping the queen by foraging for food, guarding the nest, and caring for young, they increase the offspring production of the queen by a rate of *β* bees per day per helper. (b) Illustration of one possible reproductive strategy. The phenotype of a queen is defined by four traits: (*m*_0_, *m*′, *h*_0_, *h*′). *m* is the probability that each offspring is a male. *h* is the probability that each nonmale offspring is a helper. On day 1, the values of *m* and *h* are set at *m*_0_ and *h*_0_, and these values are incremented by *m*′ and *h*′ each day. The example shown is (0.3, 0, 0.75, -0.004). (c) Haplodiploid genetics of the theoretical model. Each queen has two sets of trait-determining genes, one of two randomly determined from the mother (A,B,C,D and a,b,c,d) and one from the father (A_m_, B_m_, C_m_, D_m_). Queen trait values are the average of the two sets of alleles. Because Hymenoptera are haplodiploid, each male inherits one randomly determined set of alleles from the mother. For each new reproductive offspring, each individual allele has a probability *μ* of mutating by addition of a normally distributed perturbation with mean 0 and standard deviation *σ*. For illustration, mutated genes are marked with an asterisk.

### Evolutionary model

The behavior of each reproductive female in our model is modeled by the phenotype (*m*_0_, *m*′, *h*_0_, *h*′). This phenotype is characterized by four evolvable, heritable traits: a male fraction (*m*_0_) at the start of the season, a helper fraction (*h*_0_) at the start of the season, an increment per day (*m*′) for the male fraction as the season progresses, and an increment per day (*h*′) for the helper fraction as the season progresses. *m*_0_ and *h*_0_ are continuous and bounded between 0 (all females/reproductives) and 1 (all males/helpers); *m*′ and *h*′ are continuous and bounded between -0.01 per day (where *m* and *h* decrease by 1% each day of the season) and 0.01 per day (where *m* and *h* increase by 1% each day). This suite of alleles is inherited from one generation to the next in a haplodiploid manner (Figure 1c). For each of the four traits, females have two alleles and the overall value of that trait is the average of the values of those two alleles.

Males have only one allele for each trait. While traits are only relevant to reproductive females, the underlying alleles are also transmitted via males.

Bees with different suites of traits compete in terms of reproductive production each year. On any given day, the number of offspring of a given reproductive female is modeled by a Poisson distribution:

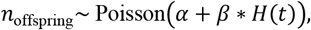

where *α* is the base average number of offspring per day, *β* is the benefit per helper, and *H*(*t*) is the total number of helpers who live in that reproductive female’s nest at time *t*. The type of each offspring is determined stochastically, with the following probabilities for a male, helper, or reproductive female at time *t*:

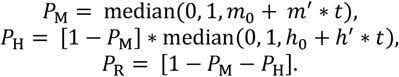

The median function ensures these probabilities are floored at 0 and capped at 1. Every haploid male offspring inherits one of the two alleles from its mother for each trait, either with no linkage disequilibrium (each gene inherited independently of the others) or, in alternative simulations, with perfect linkage disequilibrium (the four genes inherited as a single block). We only compared these two extremes, though intermediate cases are also possible. Each reproductive diploid female inherits, for each trait, one of two genes from her mother (either with or without linkage disequilibrium, as for males) and the one corresponding gene from her father (Figure 1c). After birth, each immature offspring takes (*τ*_M_, *τ*_H_, *τ*_R_) days to become a mature male, helper, or reproductive female, respectively (between 20-70 days across Hymenopterans (17)). When an individual becomes a mature male, we add him to the “pool” of always available, mature males. When an individual becomes a mature reproductive female, we add her as a new mother with initially 0 helpers. This reproductive female is also immediately fertilized by a randomly selected mature male (when *M*(*t*) ≥ 5) from the pool, which does not affect the female’s genes or traits but does affect her daughters’. To allow for evolutionary change, each individual gene in every reproductive offspring has a probability *μ* of mutating by addition of a normally distributed perturbation with mean 0 and standard deviation *σ*. The magnitude of *σ* depends on the associated trait (Table 1). When a helper matures, the average daily number of offspring per day of her mother increases by *β*. Each day, every individual has a chance (*γ*_M_, *γ*_H_, *γ*_R_) of dying and being removed from the model. The expectation of the total change in males, reproductives, and helpers for all reproductives with the same strategy on a given day is:

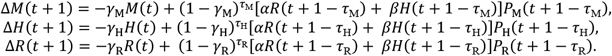

**Table 1.**
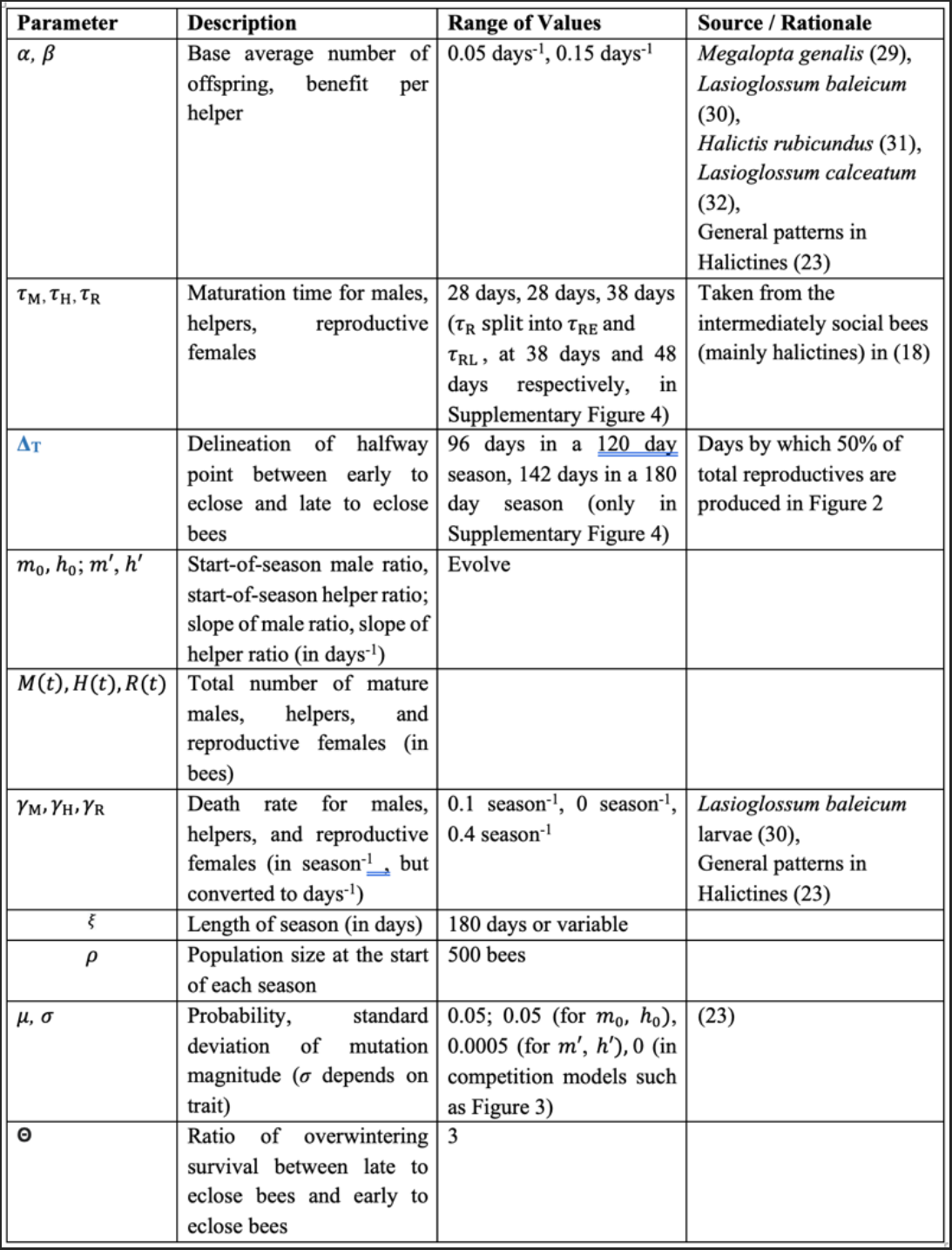
Parameters employed in our genetic model. Greek letters are intrinsic to the model, lowercase English letters evolve from season-to-season, uppercase English letters change over a season.

| Parameter | Description | Range of Values | Source / Rationale |
| --- | --- | --- | --- |
| $\alpha, \beta$ | Base average number of offspring, benefit per helper | 0.05 days<sup>-1</sup>, 0.15 days<sup>-1</sup> | *Megalopta genalis* (29), *Lasioglossum baleicum* (30), *Halictus rubicundus* (31), *Lasioglossum calceatum* (32), General patterns in Halictines (23) |
| $\tau_M, \tau_H, \tau_R$ | Maturation time for males, helpers, reproductive females | 28 days, 28 days, 38 days ($\tau_R$ split into $\tau_{RE}$ and $\tau_{RL}$, at 38 days and 48 days respectively, in Supplementary Figure 4) | Taken from the intermediately social bees (mainly halictines) in (18) |
| $\Delta_T$ | Delineation of half-way point between early to eclose and late to eclose bees | 96 days in a 120 day season, 142 days in a 180 day season (only in Supplementary Figure 4) | Days by which 50% of total reproductives are produced in Figure 2 |
| $m_0, h_0; m', h'$ | Start-of-season male ratio, start-of-season helper ratio; slope of male ratio, slope of helper ratio (in days<sup>-1</sup>) | Evolve | |
| $M(t), H(t), R(t)$ | Total number of mature males, helpers, and reproductive females (in bees) | | |
| $\gamma_M, \gamma_H, \gamma_R$ | Death rate for males, helpers, and reproductive females (in season<sup>-1</sup> but converted to days<sup>-1</sup>) | 0.1 season<sup>-1</sup>, 0 season<sup>-1</sup>, 0.4 season<sup>-1</sup> | *Lasioglossum baleicum* larvae (30), General patterns in Halictines (23) |
| $\xi$ | Length of season (in days) | 180 days or variable | |
| $\rho$ | Population size at the start of each season | 500 bees | |
| $\mu, \sigma$ | Probability, standard deviation of mutation magnitude ($\sigma$ depends on trait) | 0.05; 0.05 (for $m_0, h_0$), 0.0005 (for $m', h'$), 0 (in competition models such as Figure 3) | (23) |
| $\Theta$ | Ratio of overwintering survival between late to eclose bees and early to eclose bees | 3 | |

We obtained solutions of these equations both deterministically and via stochastic simulations.

We ran the simulation for *ξ* days. After that timepoint, we randomly selected enough mature reproductive females without replacement so that each year starts with *ρ* reproductive females, and we removed all helpers and males.

### Estimating Season Length in Regions of Social Polymorphism

To contextualize our model, we sought to compare the conditions associated with social and solitary outcomes in the model to the climatic conditions associated with social versus solitary nesting in two well-characterized sweat bee species. First, we examined the season lengths of the social and solitary populations of *Halictus rubicundus*. This species occurs both in Europe and in North America; it is typically eusocial, but solitary populations have been documented at high elevations in the United States (4) and at high latitudes in the UK and Scotland (2, 31). We also examined the locations of social and solitary populations of a second, socially polymorphic sweat bee, *Lasioglossum calceatum* (32). This species is distributed throughout the palearctic, and it is most commonly eusocial (33). Solitary populations of *L. calceatum* have been documented in the northern UK and Ireland (32) as well as at high elevations in Hokkaido, Japan (3).

To estimate season lengths for the solitary and social populations of each of these species we found data available for all locations for the period 1965 to 1975 using the Center for Environmental Data Analysis (CEDA) and the Climate Data Online Search (CDOS) for England and the US, respectively (44, https://www.nrcs.usda.gov/wps/portal/wcc/home/). We elected to limit our data to these years in order to minimize the impacts of climate change. Some of our data contained many missing days. In order to allow us to use this data, we generated 100 uncorrelated seasons from this data using RMAWGEN (https://rdrr.io/cran/RMAWGEN/man/RMAWGEN-package.html, version 1). This also reduced the impact of local weather conditions and correlation between days. We used the generated weather data to estimate the season length for each of these locations by calculating the length of time between the end of the last day of the first 5-day interval where the max temperature each day was above 14°C, approximately the minimum temperature required for *Halictus rubicundus* and *Lasioglossum calceatum* to forage (3,13) and the end of the last day of the first 5-day interval after that where the max temperature each day was below 14°C.

## Results

### Emergence of eusocial and solitary reproductive strategies

We observed the emergence of both solitary and eusocial reproductive strategies that are consistent with previous studies (Figure 1; 17, 23). In lineages with a solitary strategy, founding females tend to produce a mix of reproductive males and females at the outset of the season. Reproductive females mate with males from other nests, found their own nests later that season, and produce their own reproductive males and females (4, 17, 27, 34-35). In social lineages, founding females produce nonreproductive females early in the season, echoing the sex-ratio skews previously described in Seger’s deterministic model for the evolution of sex ratios in bivoltine species (17). The nonreproductive workers do not produce offspring of their own, but can instead increase the number of offspring produced by their reproductive mother through foraging, guarding, and nursing (23).

Our model stabilized at two equilibria analogous to these behavioral strategies, one with a high *h*_0_ and one with a low *h*_0_ (Figure 2). In the first equilibrium (the “solitary strategy”), all nests produce males at the start of the season along with a smaller number of reproductives and helpers (this value is not exactly 0 because the range for *m*_0_ and *h*_0_ is 0-1, and as such mutations will always move the values away from the extrema). Later in the season, more reproductive females and fewer males are produced (Figure 2a’). Effectively no helpers are produced. In the second equilibrium (the “social strategy”), all nests produce helpers at the start of the season; at the end of the season, a larger number of reproductive females and males are produced (Figure 2b’, Supplementary Figure 1). If *β* is decreased to 0, the social strategy disappears entirely (Supplementary Figure 2). We compared stochastic simulations of these two equilibria (error bars in Figure 2a’ and 2b’) with the corresponding deterministic solutions (dashed curves) over the course of a single season, with extremely high consistency between the two.

**Fig. 2.**
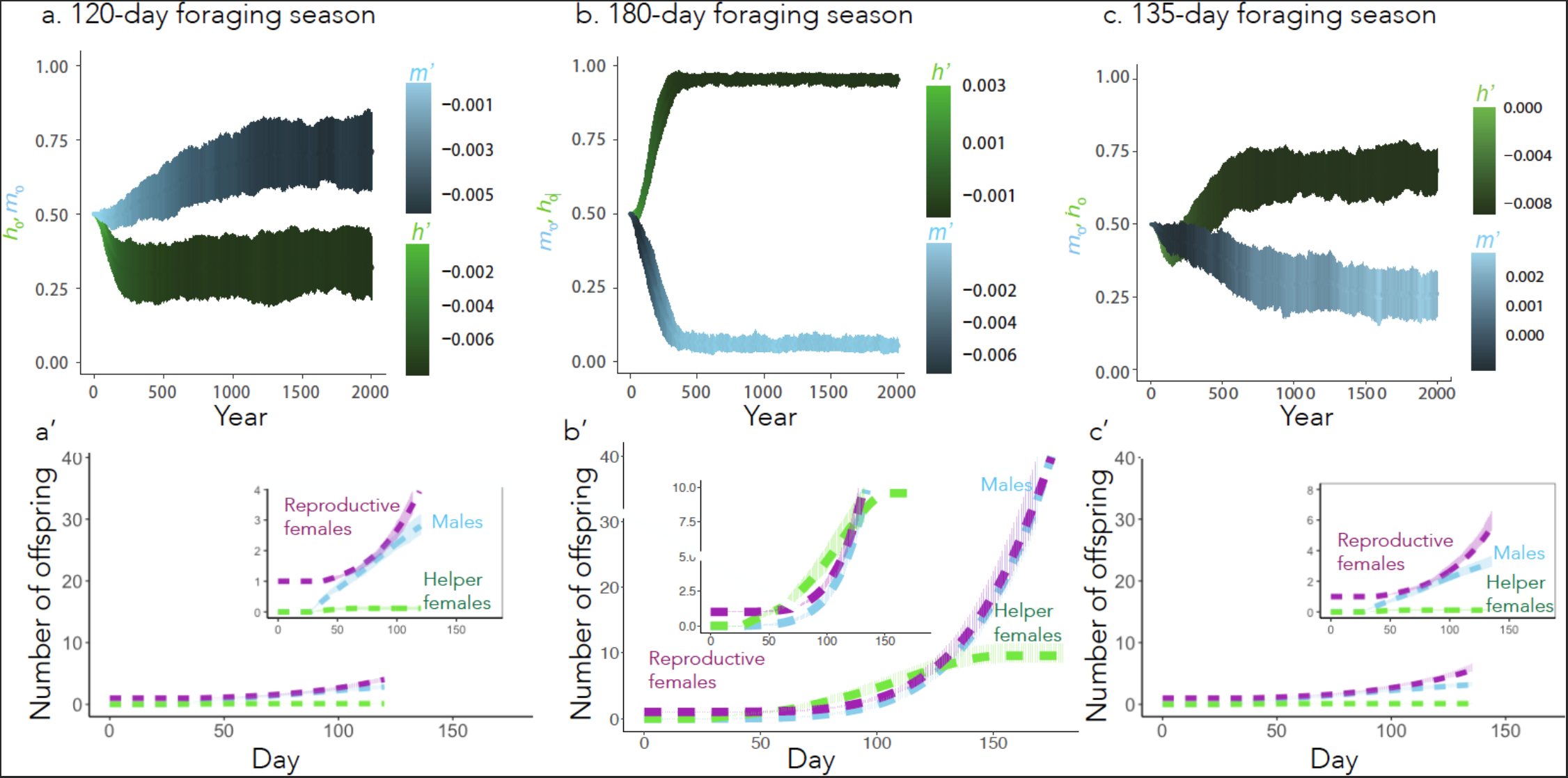
Both solitary and social equilibria can emerge, depending on length of the foraging season. (a-c) Emergence of alternative behavioral strategies over 2000 years. The starting values and for each year are represented by the centroids of the blue and green bars, respectively, while the mean slopes and are depicted in color. Bars represent the mean ± SD of 100 simulations. The season length is 120 days in (a), 180 days in (b), and 135 days in (c). (a’-c’) Cumulative number of each class of offspring over a season derived from a single initial queen with the mean strategy at year 2000. The dashed curves represent the deterministic model (Eqs. 1-3) and the pale bars represent the mean ± SD of the 100 stochastic simulations. Insets in these figures are zoom ins of the main panels for clarity.

Our simulations revealed that an approximately equal total number of males and reproductive females were produced by the end of their respective seasons (about 3 males and females per queen in Figure 2a’, and about 40 males and females per queen in Figure 2b’), though there is variation in when in the season males/females are produced. This suggests that the strategies are well optimized for the end of the year, as a 1:1 sex ratio is typically the most evolutionarily stable strategy in simulations (36). However, in natural systems, social insects often produce more females than males (37). In our simulation, we assume that workers benefit the production of males and females equally, which may not be the case in observed natural systems.

### Longer foraging seasons favor social strategies

Season length strongly influences the emergence of the two behavioral equilibria. At a sufficiently short season length (*ξ* = 120 days), the phenotype approaches (0.71, −0.0055, 0.32, −0.0078) (Figure 2a), in which males are produced early in the season followed by reproductive females (Figure 2a’). At a sufficiently long season length (*ξ* = 180 days), a phenotype emerges (0.054, 0.0031, 0.95, −0.0075), in which helpers are produced early in the season followed by reproductive females and males (Figure 2b’). At intermediate season lengths (*ξ* = 135 days), an intermediately social equilibrium emerges (Figure 2c and 2c’), which produces fewer helpers and more males early in the season relative to the 180-day social strategy.

Relatedly, helper production is disfavored at a season length of *ξ* = 180 days when the maturation time of all individuals was increased (*τ*_M_, *τ*_H_ *τ*_R_ = 68, 68, 78 days – approximately the average development time for solitary bees in (18)) (Supplementary Figure 3). This is in accord with past work showing that the interplay between season length and development time can have important implications for the life-history strategies of bee societies (18). It is worth noting that the maturation time of reproductives is not always consistent across a year. Many bee species produce winter-destined reproductives later in the year, which have a longer development time, but are better able to survive in the winter (16). To incorporate this fact, we added a parameter in the model to change the maturation time for bees before the midpoint of the foraging season. We then added a parameter Θ to represent the increase in probability of an overwintering-destined bee successfully overwintering. These changes had little effect on the relative fitness of social or solitary morphs during long season lengths (Supplementary Figure 4b), but did increase the value of *h*_0_ when the season length was short (Supplementary Figure 4a).

### Season length underlies competition and coexistence among strategies

We next evaluated the ability of each strategy to successfully invade when the other strategy was dominant. We initiated the simulations with all reproductive females at one equilibrium (social or solitary), and after 200 years, a fraction (10%) of reproductive females with the other strategy immigrates into the population. We lowered the value of *µ* to 0 (Table 1) so that only immigration could drive phenotypic change. If all nests are initially social, solitary foundresses are only able to successfully invade when the season length is below 135 days (Figure 3a-b). If all nests are initially solitary, social foundresses are only able to successfully invade when the season length is above 155 days (Figure 3d). In between these two limits is a region where both strategies are resistant to complete removal by the other (Figure 3c). This results in the emergence of an intermediate region (135-155 days) where both strategies can coexist.

**Fig. 3.**
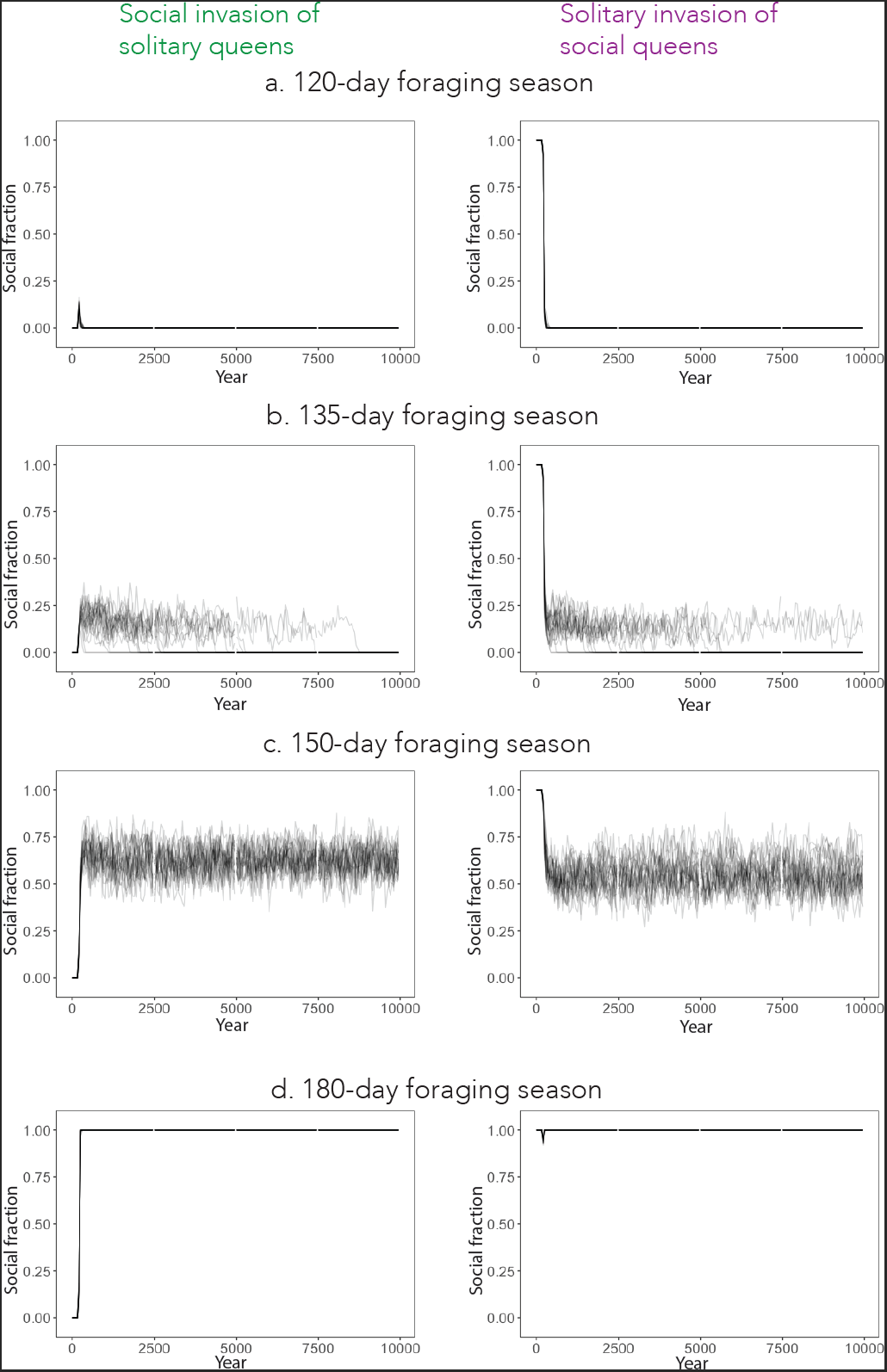
Simulated invasions reveal regions of invasibility for social and solitary strategies. We began simulations in an environment either with 100% solitary queens (0.71, -0.0055, 0.32 -0.0078), on the left, or 100% social queens (0.054, 0.0031, 0.95, -0.0075), on the right. After 200 years, we replaced 10% of queens with the opposite strategy (social on the left, solitary on the right). We estimated the social fraction from the mean value of h_0_ for each reproductive female that survives to the end of the season, using that to determine which bees were social/solitary. Each plot overlays 25 independent simulations.

In these simulations, the four traits that define the behavioral phenotype are determined by unlinked loci. As such, defining the fraction of social queens in the population based on *h*_0_ alone only reveals part of the story. We therefore also considered the fraction of social queens based on the other three loci for a 150-day foraging season. Intriguingly, while *h’* and *m’* both eventually fixed to all solitary and all social values respectively, both the social and solitary values of *h*_0_ and *m*_0_ coexisted in the population (Supplementary Figure 5), though at somewhat different ratios. While presence of the social allele of and the social allele of were correlated significantly (*p* < 2E-16), the correlation is very weak (*r* = - 0.059), suggesting a split sex ratio where more social species are also more likely to be female biased. The low correlation makes sense because our four alleles frequently reassort in our simulation without linkage disequilibrium.

Because our alleles reassort after every generation, the F1 hybrid generation of reproductives that emerge after an invasion might not be able to successfully optimize male and helper ratios and would therefore be less fit. This would thus favor the behavioral strategy which is already dominant in the population. Similar reductions in hybrid fitness have been seen in a range of bee species (38-39). This is consistent with Supplementary Figure 5, as a hybrid won’t be able to optimize its male and helper fractions together. We therefore considered what would happen if all four trait-determining loci were inherited as a linked block. We simulated this case of maximal linkage disequilibrium at 154 days, as at that season length the simulation fixes to the social population, but very slowly. While linkage disequilibrium didn’t result in the social population invading faster, it was significantly associated with the maintenance of coexistence of both behavioral forms (Supplementary Figure 6, χ^2^ = 18.9, *p* < 1E-5).

Coexistence can often be dependent on the parameters of a model (40). We implemented two substantial changes to parameters to see if they changed the emergence of a coexistent regime: decreased benefit per helper or increased male mortality (Supplementary Figure 7). Neither change removed the ability of these two strategies to coexist, though decreasing the benefit per helper shifted the range of season lengths where coexistence occurs. In principle, the intermediate region of coexistence might disappear or become larger if the parameters of the model are changed in other ways, but we do not explore this further here.

### Model outcomes mirror natural populations of halictine bees

To assess the translational potential of our model, we compared our theoretical transitional boundaries to known examples of solitary-social dimorphisms in a group of socially variable sweat bees (Hymenoptera: Halictidae). In this family of bees, some species are socially polymorphic, and exhibit eusocial or solitary behavior in different parts of their range (26, 41). First, we looked at transition zones in behavior for *Halictus rubicundus*, a sweat bee species known to be socially polymorphic in both North America (34) and the United Kingdom (2). In North America, *H. rubicundus* lives in social nests in most of Colorado, but in the higher elevation around Crested Butte, Colorado, *H. rubicundus* exhibits a solitary phenotype (4). We generated weather data for both Crested Butte and the neighboring town of Almont from empirical weather data to estimate the difference in season length between the two locales (Supplementary Table 1).

Interestingly, we found that the solitary populations of *H. rubicundus* in Crested Butte experience season lengths that fall below the intermediate region of the model (at approximately 120 days) while the neighboring, social populations in Almont experience season lengths that fall above the upper limit of the intermediate region (at around 154 days) (Figure 4a). This suggests that the predictions made by the model can explain the patterns of social and solitary strategies in a natural environment.

**Fig. 4.**
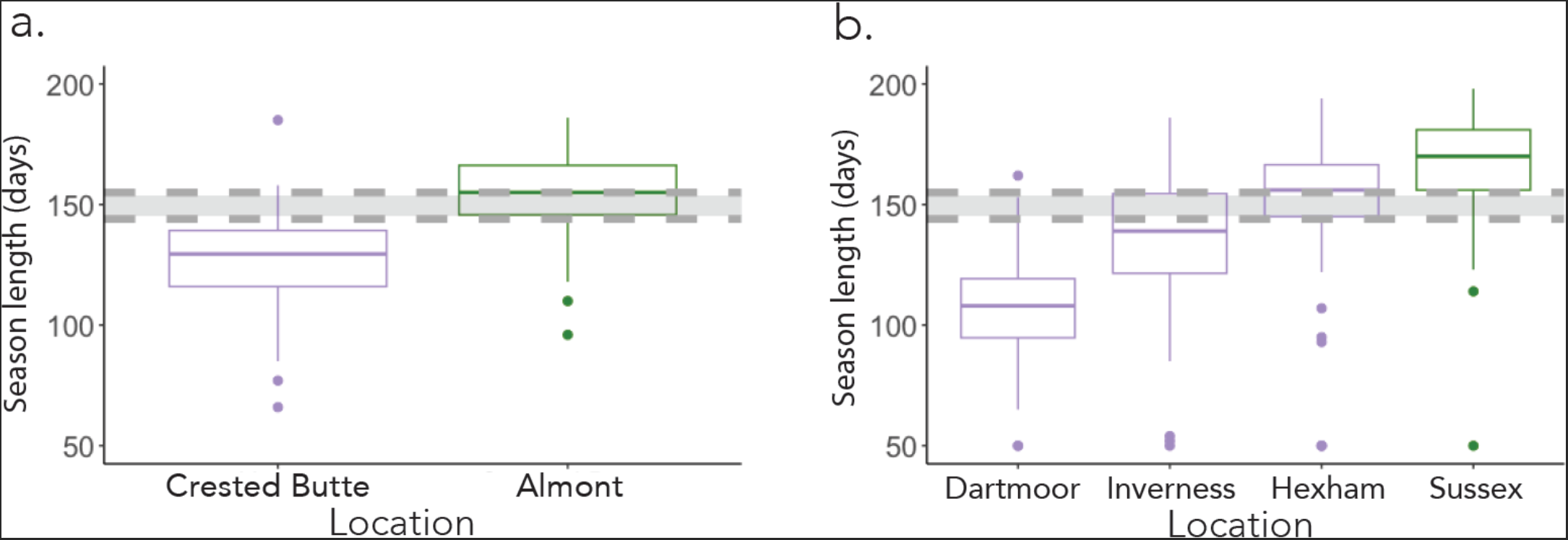
Model predictions are well matched to behaviors observed in natural populations of socially variable bees. Weather data was obtained from the CEDA and CDOS (see Methods) and then generated using RMAWGEN, and season length was calculated as the time between the end of the last day of the first 5-day period where the temperature each day was above 14°C and the end of the last day of the first 5-day period after that where the temperature each day was below 14°C. The box-and-whisker plot shows the interquartile range. Colors represent the social behavior of *Halictus rubicundus* (a) or *Lasioglossum calceatum* (b), where purple is solitary, and green is social. The dark grey region in each figure represents the region of coexistence identified in Figure 3.

We similarly analyzed data on *Lasioglossum calceatum*, a sweat bee that varies in social behavior across the United Kingdom. We considered four regions in the United Kingdom where *L. calceatum* exhibits variant behavior: social behavior in Sussex, and solitary behavior in Inverness, Hexham, and Dartmoor (32,42) According to our generated weather data Sussex has a mean season length that falls above the intermediate region, at around 175 days, while in Dartmoor and Inverness, where *L. calceatum* produces solitary nests, mean season lengths fall within or below the intermediate region, between 110 and 150 days. Hexham is the exception to this, having a mean season length that falls slightly above the intermediate region (Figure 4b). This may be due to other geographic factors, such as a higher windspeed in Hexham compared to the other cities (43-44). Taken in sum, this suggests that (i) our model does an accurate job of predicting patterns of social versus solitary behavior, and (ii) there is frequently variation in social behavior in natural populations where season lengths fall within or near the intermediate region we identify in our model.

## Discussion

### Social and solitary behavioral patterns emerge in a freely evolving system

We developed an individual-centered model that allows several life history traits to vary freely. This model expands upon two pivotal theoretical models (17, 23) by incorporating two additional variables: season length and offspring development time. The incorporation of these variables allows us to extend these existing frameworks to study how the environment impacts the emergence and evolution of eusociality and how the costs and benefits of social living vary across environmental gradients.

Seger developed one of the first models to show that biased sex ratios emerge in species with a partially bivoltine life history. Quiñones and Pen built on this model by looking not just at the emergence of biased sex ratios, but how helping behavior emerges. Again, using a partially bivoltine model, they identified a confluence of individual preadaptations which are necessary conditions for the emergence of helping behaviors and split sex ratio emblematic of many social insects.

Our results demonstrate that by simply manipulating season length and offspring development times, our model can closely match patterns of life history and social variation observed in nature. Although we did not explicitly include voltinism (i.e. discrete broods) or sex ratios as variables in our model, we found that both evolve as emergent properties in this system. For example, we observed that, within our model, the solitary strategy produced males first, followed by a steady increase in the production of reproductive females (17, 37). Similarly, the social strategy produced helpers followed by a steady increase in the production of both reproductive males and females (26).

### Different season lengths favor different behavioral equilibria

Our model underscores the importance of incorporating environmental parameters into theoretical models. We found that a social equilibrium emerged when foraging seasons were long and egg-to-adult development times were relatively short (Figure 2, Supplementary Figure 2). This is consistent with assured fitness returns models (45-46) that associate the selective value of a helper with the length of time she can provide a “return” on investment relative to producing a reproductive. If a population either cannot get enough benefit from helpers because the season is too short, or because it takes too long for helpers to mature and begin helping, a strategy based around helping stops being effective. By the same token, work in game theory has shown that short-term investments with long-term gains are favored when there is more time to reap those gains (47-49). Our work provides a quantitative framework that allows us to generate hypotheses and test predictions about how season length shapes social behavior and its evolution.

### Social and solitary strategies coexist at intermediate season lengths

We next tested the ability of social and solitary strategies to invade an existing population that employs the opposite strategy. Below a certain season length (135 days in the model), the solitary strategy always took over, regardless of starting conditions. Above a certain season length (155 days in the model), the social strategy always took over. Between those two season lengths, however, neither strategy could totally outcompete the other, and both coexisted in the model (Figure 3b). We believe that, in this region, the two strategies are likely to be maintained by balancing selection. In an environment where all bees are social, solitary males have less competition, so even if the social strategy more efficiently produces females, they will be disproportionately fertilized by males from solitary colonies. Moreover, in an environment where all bees are solitary, there are diminishing returns to producing males early in the season because of the substantial competition among them. We did not find changes in parameters that lead to a loss of this coexistence regime (Supplementary Figure 7), though decreasing the helper benefit *β* increased the minimum season length required for the social strategy to emerge.

The presence of coexistence provides a theoretical framework for understanding the presence of behavioral polymorphism within one population. There are striking examples in nature of social and solitary phenotypes of the same species coexisting in one population. For example, *Lasioglossum baleicum* females (50) produce both eusocial and solitary nests within a single, panmictic population in Hokkaido, Japan. In this population, differences in soil temperatures and sunlight are highly correlated with this variation, potentially providing a fascinating empirical system to further test our model predictions. Such studies can lead to a greater understanding of the evolutionary benefits of social and solitary behavioral strategies in panmictic populations.

### Linkage among traits helps drive the emergence of eusociality

Even though all four loci were entirely unlinked in our initial simulations, we still found a weak, but significant, association between the social alleles (Supplementary Figure 5). We believe that this is because both the social and solitary strategies depend on joint optimization between the male fraction and the helper fraction. This is further suggested by the fact that linking the four traits together strengthens the ability of the social and solitary alleles to coexist (Supplementary Figure 6). Many known genetic transitions between social and solitary species in nature are mediated by analogously structured “supergenes” – sets of genes which are inherited as a genetically-linked block (51) For instance, some transitions among social forms in ants are known to be associated with inversions that suppress recombination and produce supergenes with different traits inherited as a single unit (6, 52-53). To be clear, our model is far more simplistic than the supergenes found in nature; while two linked supergenes in ants are associated with colony sex ratio and social form (54), no single gene within a supergene has been found to be associated with male ratio nor helper ratio. Rather, our model is intended to provide a conceptual framework within which to understand these results.

### Environment shapes social behavior and its evolution

A growing body of evidence demonstrates the importance of environmental factors to the expression of social behavior (1,5-9, 55). For example, gradients in sociality are linked to altitudinal and latitudinal clines (reviewed in (6)) and transplantation experiments and surveys of social and solitary insect species have shown that the social composition of insect societies can respond dynamically to local temperatures and environmental conditions (2, 56). For instance, the halictine bee, *Halictus rubicundus*, shifts from a solitary to a eusocial strategy when nests are transplanted from high to low latitudes with a greater number of foraging days in the season (2). In many cases, these patterns are consistent with those observed in our model – lower altitudes and more equatorial latitudes are associated with longer seasons and increased levels of sociality. For example, socially polymorphic bees and wasps do not typically produce workers at high altitudes and latitudes (i.e. they exhibit a solitary life-history) (e.g., *H. rubicundus* (31, 34, 57); *A. aurata* (58); *L. baleicum* (50, 59); *L. calceatum* (3)). Our model clearly identifies an increase in season length as sufficient to trigger the transition from solitary to social living.

It is, however, important to note that not all social insect species display decreasing social complexity with decreasing season lengths. Organisms with obligate forms of sociality or those that live in large colonies (e.g. those that are beyond Wilson’s ‘point of no return’ and do not have the ability to live and reproduce solitarily) often show an increase in social complexity with decreasing season length (reviewed in (6, 18, 60)).

The model we present here explicitly focuses on the emergence of eusociality and on transitions between solitary and simple social reproductive strategies. Future work is needed to explore if and how factors such as season length are likely to impact the behavior of organisms in larger, more complex societies. Moreover, models incorporating behavioral plasticity alongside evolutionary change could be highly informative, especially given that many bee species are highly plastic in their social behavior (2, 61). Finally, recent work has been done to predict foraging season length in honeybee colonies, which could improve upon our empirical tools to measure season length (62).

### Predicting behavioral change in a rapidly changing climate

In view of global warming and climate change, it is essential to develop tools that can predict how species’ behavioral patterns may change as seasons get longer, warmer, and more variable (13, 55, 63) Our model explicitly incorporates season length when considering the dynamics of social evolution, demonstrating that season length is a major factor shaping the evolution of pollinator communities. Our results are in accord with observations made in nature which support the assertion that we may see an increase in the number of social nests in socially polymorphic clades of bees and a subsequent decrease in solitary strategies as global temperatures rise (13). Our model suggests that regions with intermediate season lengths can support multiple, alternate evolutionary stable states, but that as regions transition outside of these intermediate season lengths, it is likely that longer seasons will exclusively favor social forms. Moreover, other species, including mice (64), danio fish (65), and guppies (66) adjust their social behaviors in response to temperature. While those species are not eusocial, this paper provides a “proof-of-principle” for developing quantitative, theoretical frameworks to examine how and why social behaviors can emerge more readily in different ecological contexts.

Behaviors are, by definition, one mechanism that organisms can use to quickly respond to environmental stimuli. How organisms respond to new environmental stressors can change both over the course of an individual’s lifetime and across ecological and evolutionary time scales. As the effects of climate change become more pronounced, it is crucial to create more models that link individual behavioral changes with the environmental factors that underlie these behaviors.

## End Matter

### Author Contributions and Notes

DMR, NSW, SAL, and SDK designed the study. DMR and NSW wrote the models and DMR analyzed the data. DMR and SDK wrote the first draft, and all authors commented and helped to revise the manuscript. The authors declare no conflict of interest.

## Acknowledgments

We thank C. Saad-Roy, S. Wolf, and members of the Kocher, Wingreen, and Levin labs for their comments and feedback on this project. We also acknowledge funding from the National Science Foundation *through the Center for the Physics of Biological Function (PHY-1734030)* and NSF GRFP 2021276634 to DMR, NSF DEB 1754476 and NIH 1DP2GM137424-01 to SDK, and additional funding from the Lewis-Sigler Institute for Integrative Genomics. SDK is an HHMI Freeman Hrabowski Scholar.

## Supplementary Materials

**Supplementary Figure 1.**
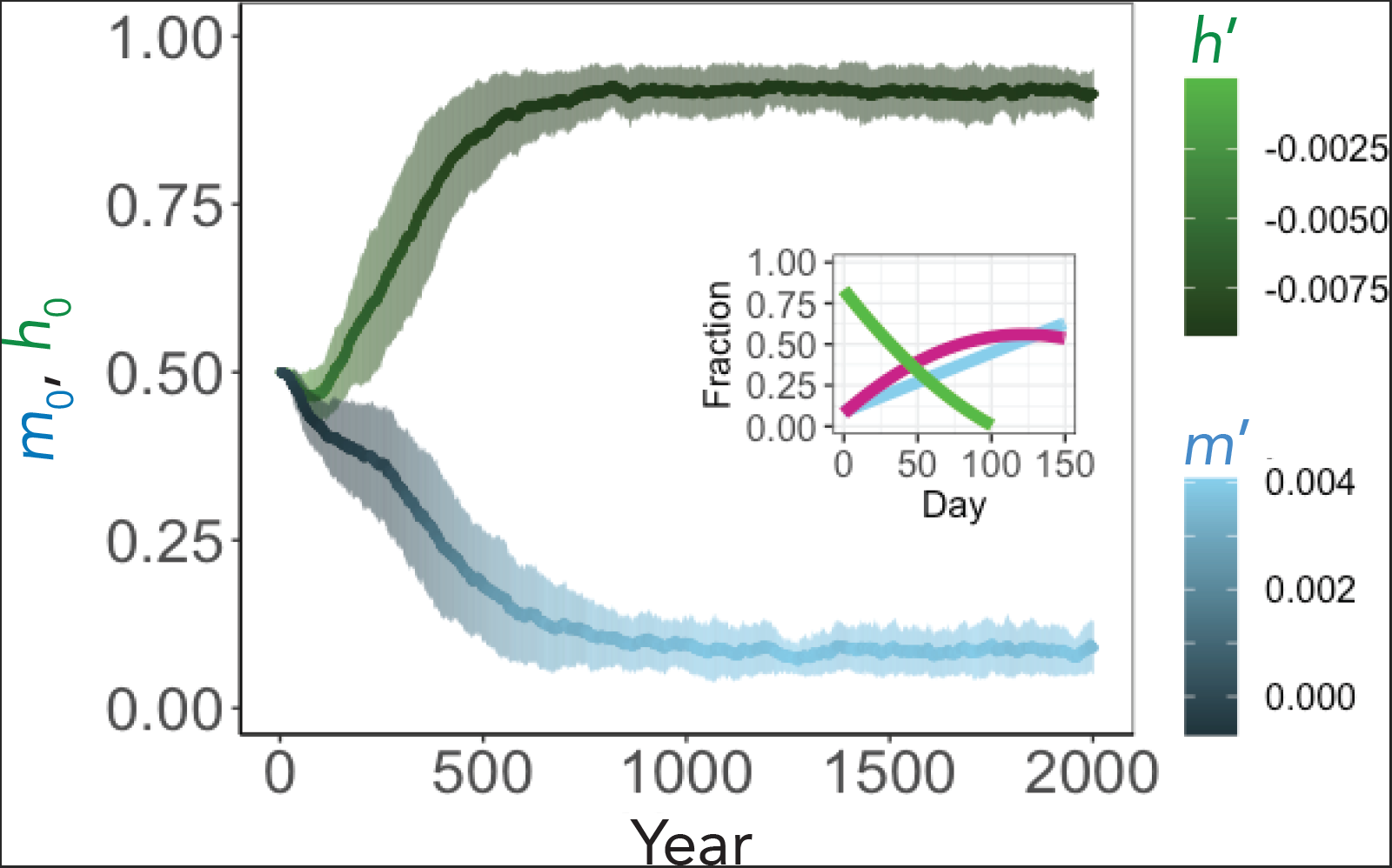
The social equilibrium emerges at 150 days. The starting values *m*_0_ and *h*_0_ for each year are represented by the centroids of the blue and green bars, respectively, while the mean slopes *m*′ and *h*′are depicted in color. Bars represent the mean ± SD of 100 simulations.

**Supplementary Figure 2.**
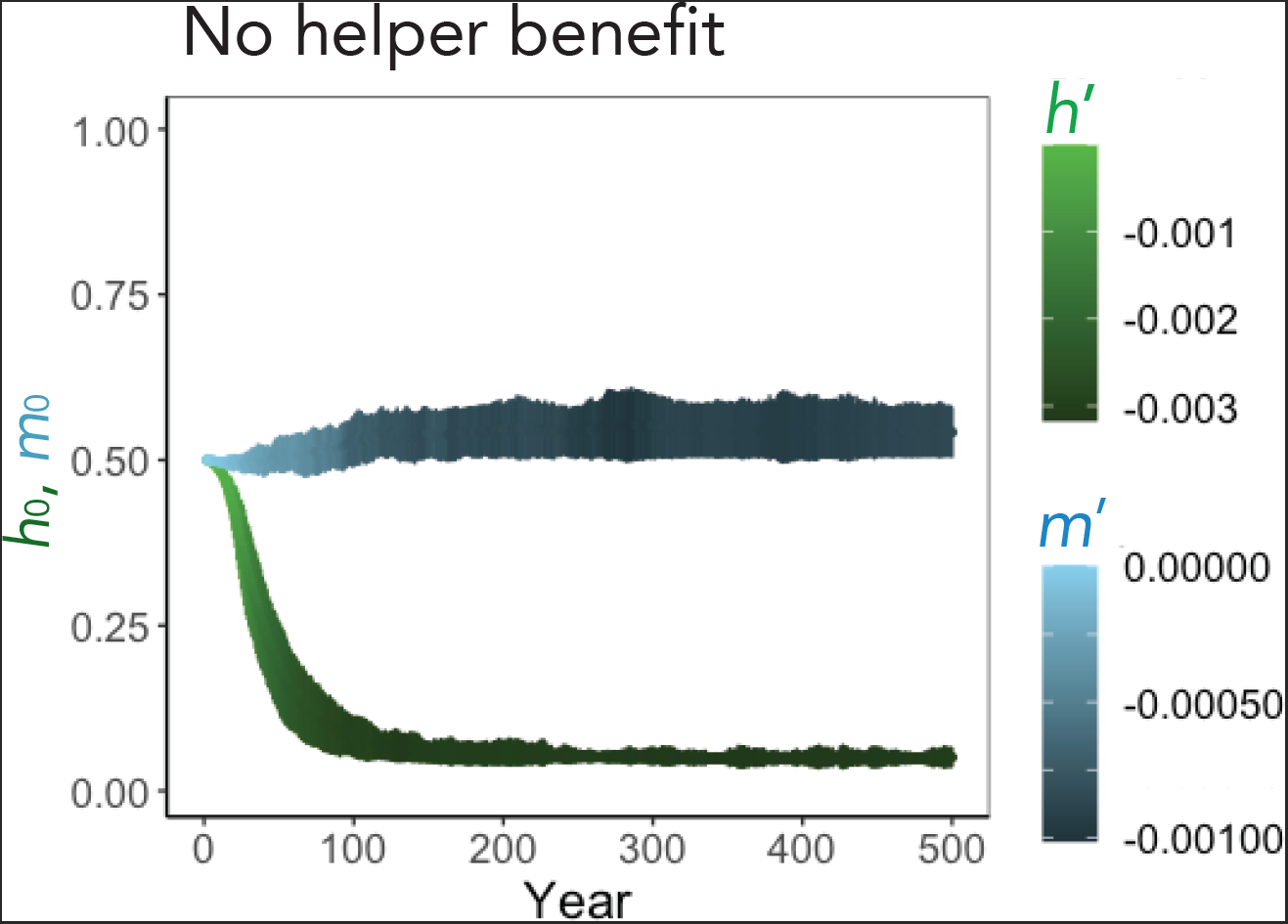
The social equilibrium does not emerge when the helper benefit *β* = 0. The starting values *m*_0_ and *h*_0_ for each year are represented by the centroids of the blue and green bars, respectively, while the mean slopes *m*′ and *h*′are depicted in color. Bars represent the mean ± SD of 100 simulations.

**Supplementary Figure 3.**
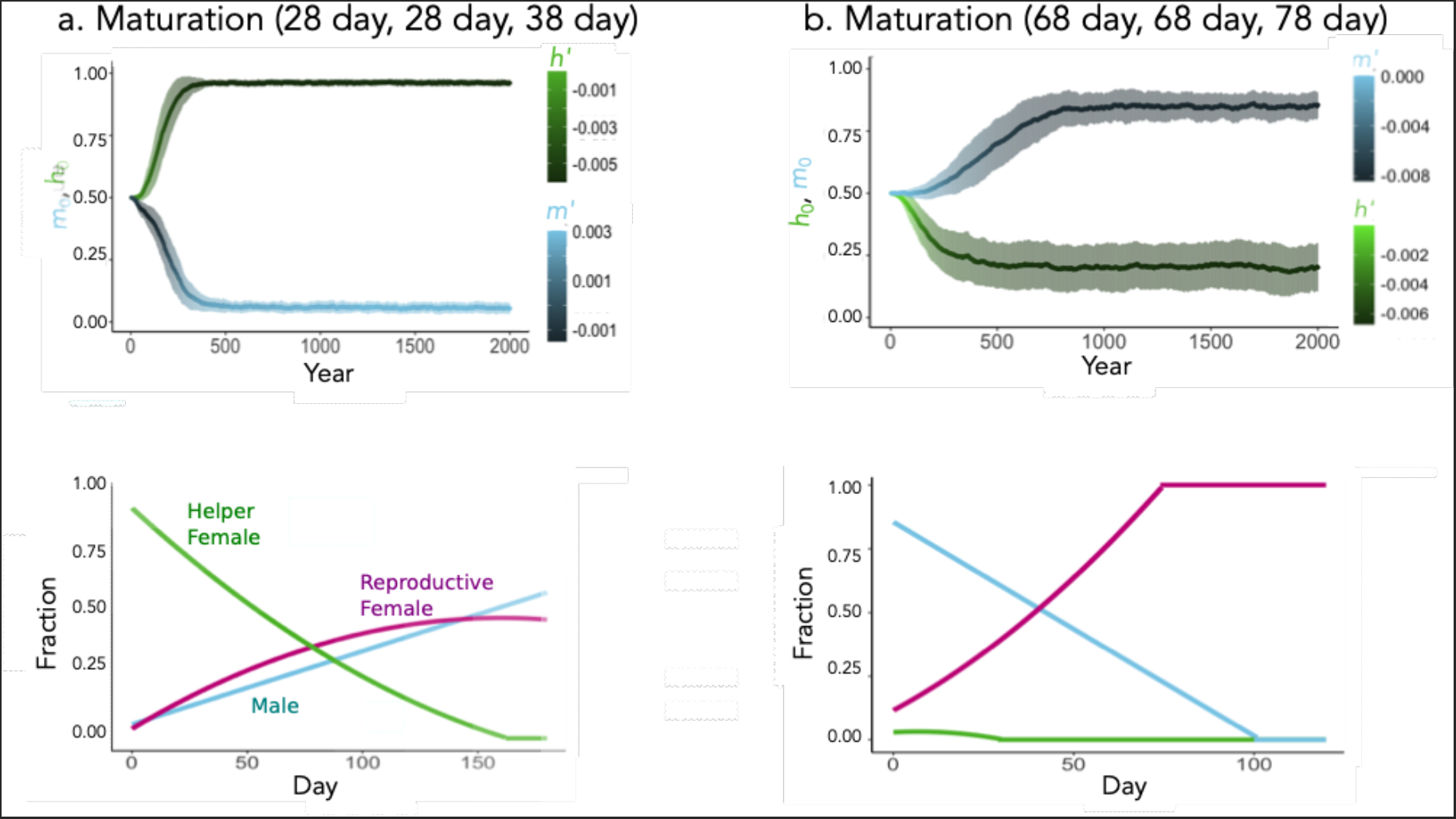
The emergence of sociality depends on maturation time. The foraging season is 180 days and the parameters are the same as in Figure 2, except that the maturation times for reproductive *males*, helpers, and reproductive females are: *τ*_*M*_ = 28 days, *τ*_*H*_ = 28 days, *τ*_*R*_ = 38 days in (a) and *τ*_*M*_ = 68 days, *τ*_*H*_ = 68 days, *τ*_*R*_ = 78 days in (b).

**Supplementary Figure 4.**
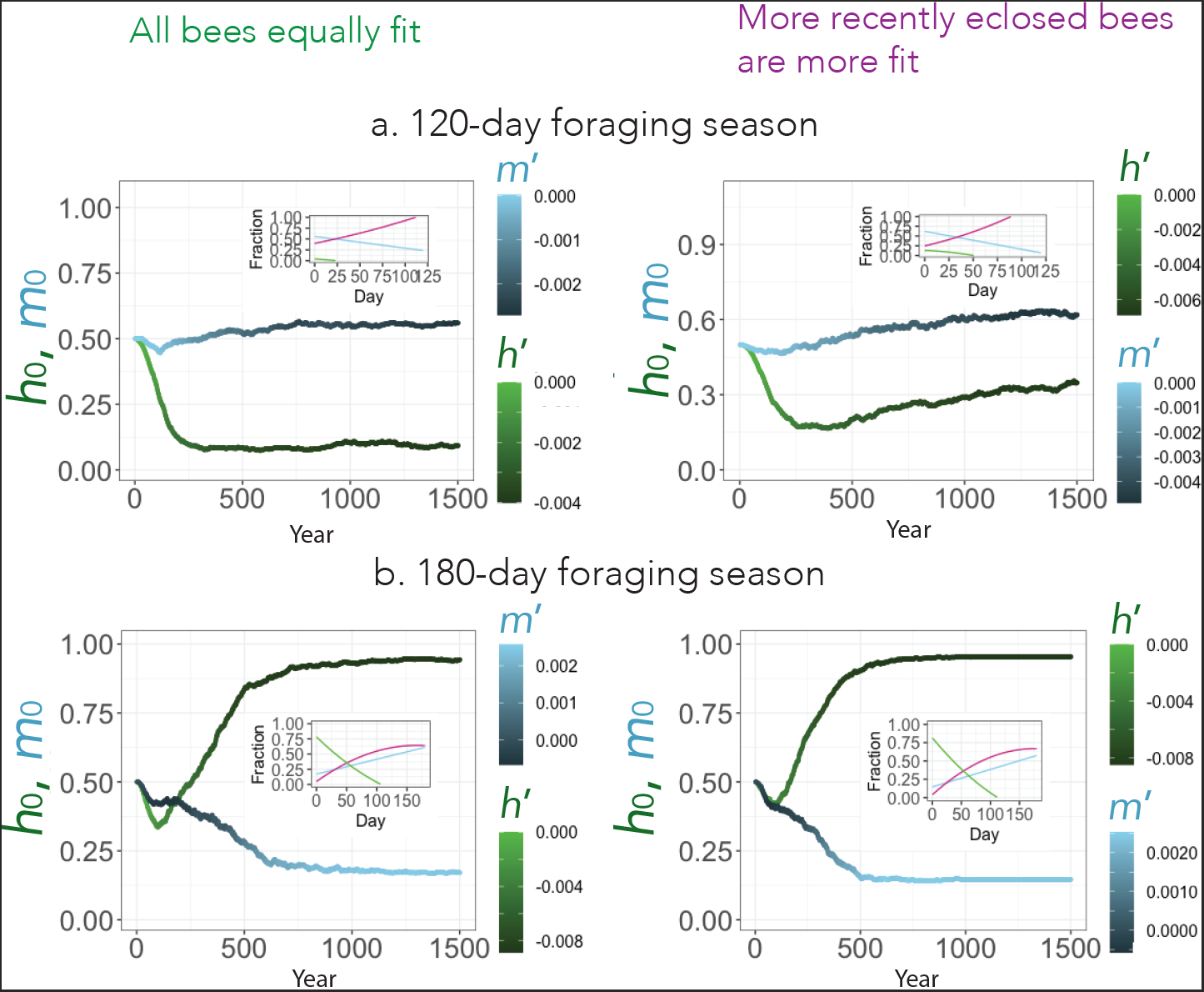
Increasing the relative fitness of late-eclosing bees that also have a longer maturation time can increase the fitness of a solitary strategy. Parameters are identical to Figure 1, with a foraging season of 120 days (a) and 180 days (b), except that the second 50% of bees to eclose (measured as bees that eclose after day 96 in (a) and after day 146 in (b)) have a *τ*_*R*_ = 48 days. Moreover, in the figures on the right, these bees also have a 3x greater likelihood of surviving the winter compared to the first 50% of bees.

**Supplementary Figure 5.**
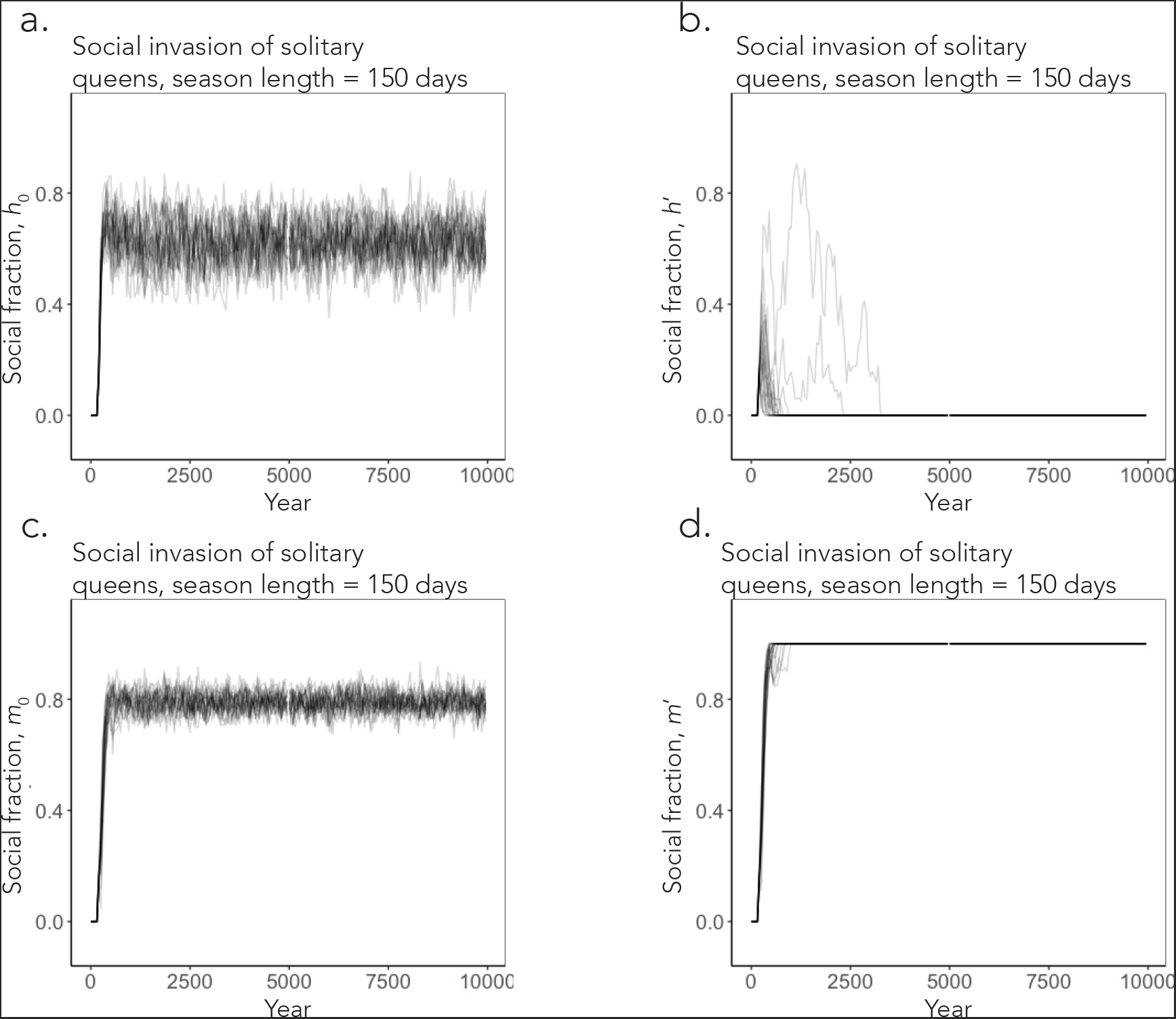
Competition between social (0.712, -0.00548, 0.318, -0.0078) and solitary (0.0537, 0.00307, 0.954, -0.00750) alleles at all four loci in our simulation for 150-day foraging season. The simulation is the same as in Figure 3c for social invasion of solitary queens (*µ* **=** 0), but with social fraction determined by the frequency of the social and solitary alleles for *h*_0_ (a), *h*′ (b), *m*_0_ (c), and *m*′ (d). Values of *h*_0_ and *m*_0_ were weakly correlated (Fisher’s exact test, odds ratio = 0.776, *p* < 2E-16, *r* = -0.059).

**Supplementary Figure 6:**
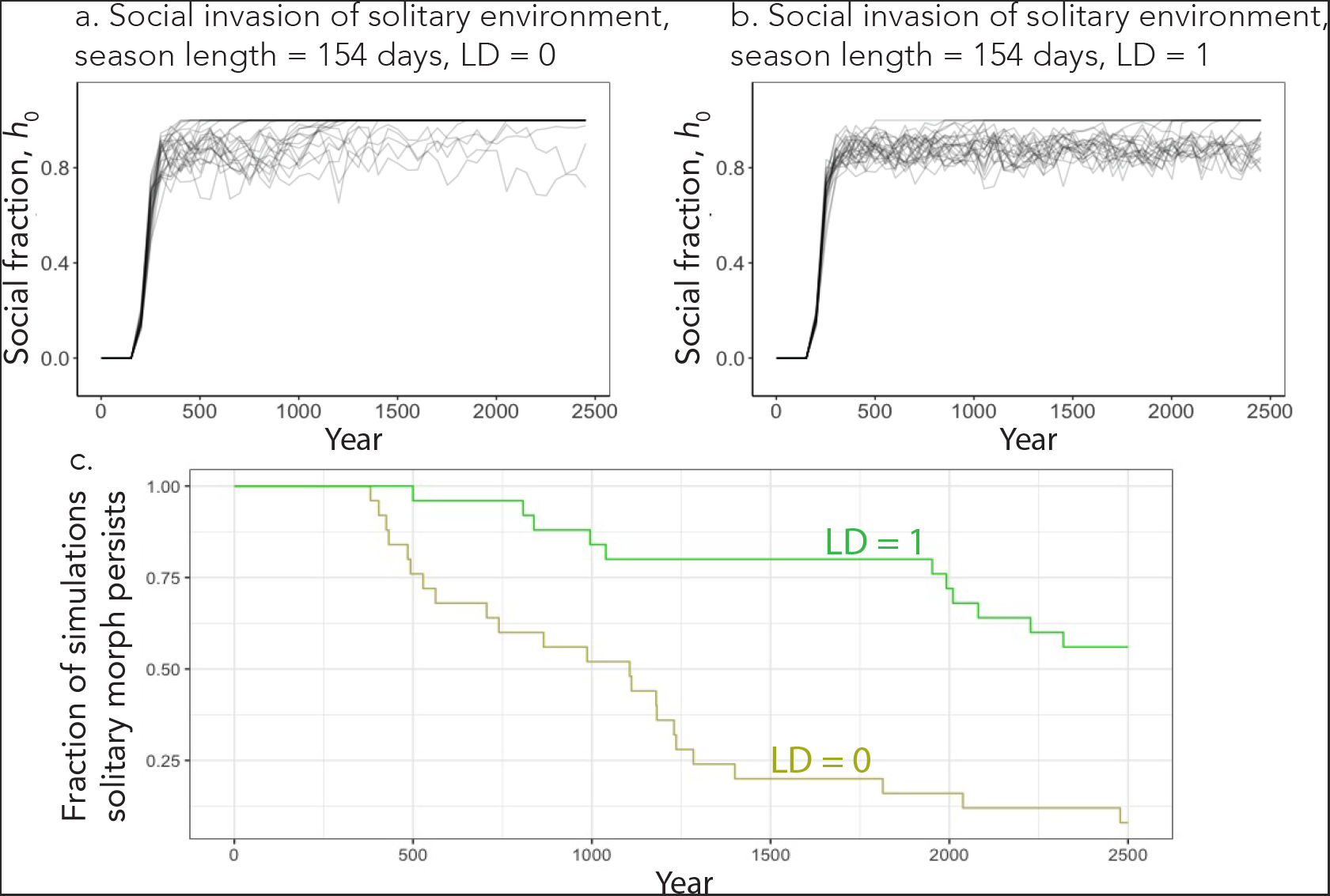
Linkage disequilibrium (LD) favors the maintenance of coexistence. The simulation is the same as in Figure 3 with a social invasion of solitary queens, except that genes are either inherited from the queen completely independently LD = 0 (a) (data from Figure 3) or as a block LD = 1 (b). (c) Persistence of the solitary phenotype for both LD = 0 and LD = 1. The *y*-axis shows the fraction of simulations where the solitary morph has not been eliminated by year 2500. The difference between the two simulations is statistically significant (rank sum test, *χ*^*2*^ = 18.9, *p* < 1.0E-5).

**Supplementary Figure 7.**
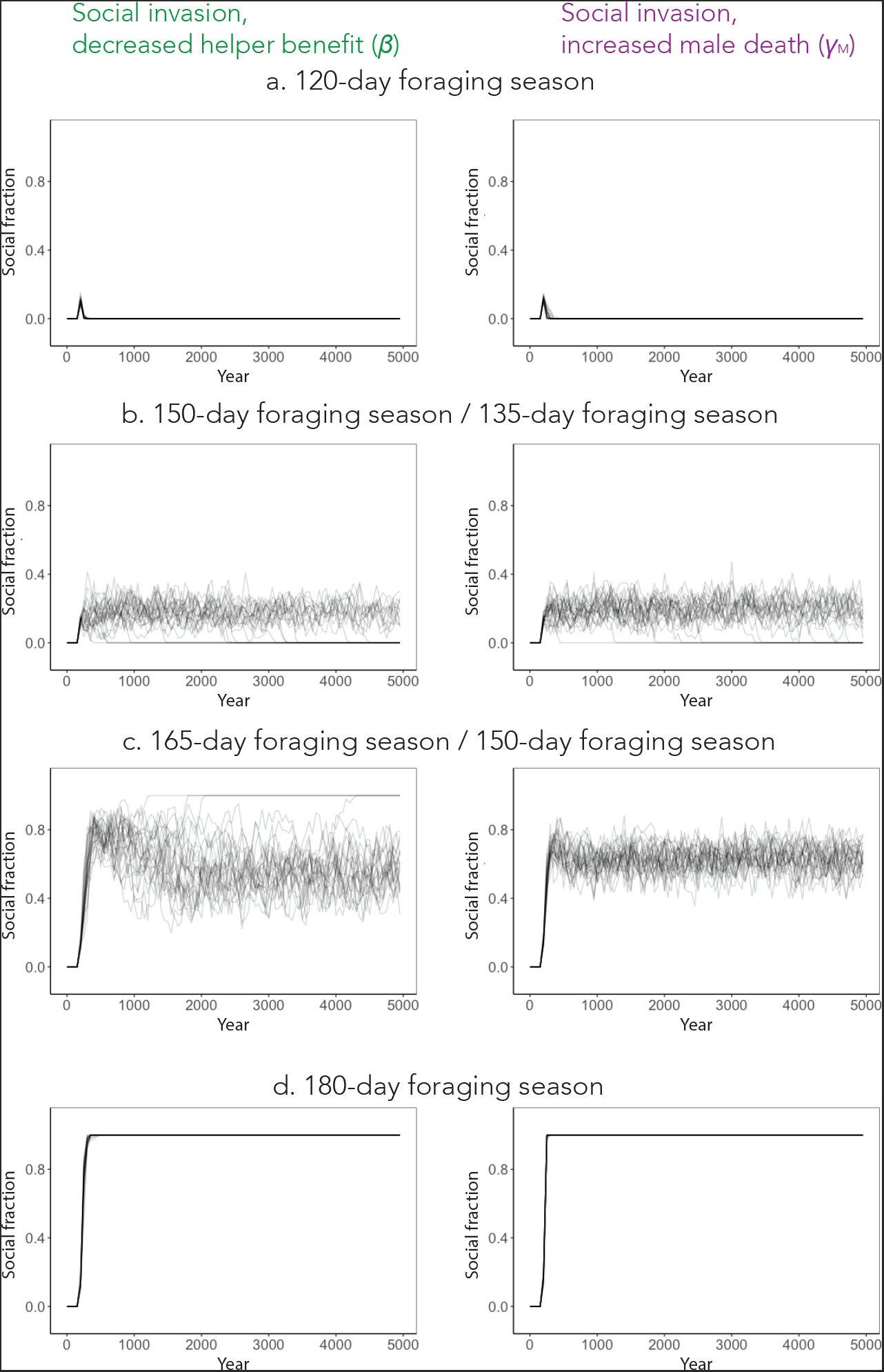
In our model, the existence of an intermediate regime of season lengths where the solitary and social strategies are mutually invasible is robust to changes in parameters. Simulations were identical to those in Figure 3b, except with decreased benefits associated with helping, *β* = 0.075 days^-1^ (a) or with an increased male death rate, *γ*_M_ = 0.01 day^-1^ (b).

**Supplementary Table 1.** Estimated season length for regions with natural variation in bee behavior. Season lengths were generated based on empirical weather measures with RMWAGEN.

| Location | Bee species | Coordinates | Behavior | 25<sup>th</sup> Quartile | Median temp. | 75<sup>th</sup> Quartile |
| --- | --- | --- | --- | --- | --- | --- |
| Crested Butte, CO | *H. rubicundus* | (38.8739° N, 106.9772° W) | Solitary | 113 days | 128 days | 140 days |
| Almont, CO | *H. rubicundus* | (38.9078° N, 106.6017° W) | Social | 144 days | 154 days | 162 days |
| Inverness | *L. calceatum* | (57.4867° N, 4.22047° W) | Solitary | 121 days | 140 days | 155 days |
| Hexham | *L. calceatum* | (54.9804° N, 2.01875° W) | Solitary | 146 days | 157 days | 167 days |
| Dartmoor | *L. calceatum* | (50.5495° N, 3.9963° W) | Solitary | 94 days | 108 days | 120 days |
| Sussex | *L. calceatum* | (50.9053° N, 0.06978° W) | Social | 160 days | 170 days | 181 days |

## References

1. Botero CA, Weissing FJ, Wright J, Rubenstein DR. Evolutionary tipping points in the capacity to adapt to environmental change. Proc. Natl. Acad. Sci. U.S.A. 2015;112(1):184–9.

2. Field J, Paxton RJ, Soro A, Bridge C. Cryptic plasticity underlies a major evolutionary transition. Curr Biol. 2010;20(22):2028–31.

3. Sakagami SF, Munakata M, editors. Distribution and Bionomics of a Transpalaearctic Eusocial Halictine Bee, Lasioglossum (Evylaeus) calceatum, in Northern Japan, with Reference to Its Solitary Life Cycle at High Altitude 1972.

4. Eickwort GC, Jeffrey ME, Gordon J, Eickwort MA. Solitary behavior in a high-altitude population of the social sweat bee Halictus rubicundus (Hymenoptera: Halictidae). Behav. Ecol. Sociobiol. 1996;38(4):227–33.

5. Lawson SP, Shell WA, Lombard SS, Rehan SM. Climatic variation across a latitudinal gradient affect phenology and group size, but not social complexity in small carpenter bees. Insect Soc. 2018;65(3):483–92.

6. Purcell J. Geographic patterns in the distribution of social systems in terrestrial arthropods. Biol Rev Camb Philos Soc. 2011;86(2):475–91.

7. Purcell J., Pellissier L., Chapuisat M. 2015 Social structure varies with elevation in an Alpine ant. Molecular Ecology 24(2), 498–507.

8. Miller SE, Bluher SE, Bell E, Cini A, Silva RCd, de Souza AR, et al. WASPnest: a worldwide assessment of social Polistine nesting behavior. Ecology. 2018;99(10):2405-.

9. Guevara J, Avilés L. Ecological predictors of spider sociality in the Americas. Glob. Ecol. Biogeogr. 2015;24(10):1181–91.

10. Purcell J, Avilés L. Smaller colonies and more solitary living mark higher elevation populations of a social spider. J Anim Ecol. 2007;76(3):590–7.

11. Liu M., Chan S.-F., Rubenstein D.R., Sun S.-J., Chen B.-F., Shen S.-F. 2020 Ecological Transitions in Grouping Benefits Explain the Paradox of Environmental Quality and Sociality. The American Naturalist 195(5), 818–832.

12. deHaan J, Maretzki J, Skandalis A, Tattersall G, Richards M. Costs and benefits of maternal nest choice: tradeoffs between brood survival and thermal stress for small carpenter bees. bioRxiv. 2022:2022.11.30.518597.

13. Schürch R, Accleton C, Field J. Consequences of a warming climate for social organisation in sweat bees. Behav. Ecol. Sociobiol. 2016;70(8):1131–9.

14. Duckworth RA. The role of behavior in evolution: a search for mechanism. Evol. Ecol. 2009;23(4):513–31

15. Renn SCP, Schumer ME. Genetic accommodation and behavioural evolution: insights from genomic studies. Anim. Behav. 2013;85(5):1012–22.

16. Hunt JH, Amdam GV. Bivoltinism as an antecedent to eusociality in the paper wasp genus Polistes. Science. 2005;308(5719):264–7.

17. Seger J. Partial bivoltinism may cause alternating sex-ratio biases that favour eusociality. Nature. 1983;301(5895):59–62.

18. Kocher SD, Pellissier L, Veller C, Purcell J, Nowak MA, Chapuisat M, et al. Transitions in social complexity along elevational gradients reveal a combined impact of season length and development time on social evolution. Proc Biol Sci. 2014;281(1787).

19. Batra SW. Nests and social behavior of halictine bees of India (Hymenoptera: Halictidae). Indian J. Entomol. 1966;28:375.

20. Szathmáry E, Smith JM. The major evolutionary transitions. Nature. 1995;374(6519):227–32.

21. Wilson EO. Sociobiology: The New Synthesis: Harvard University Press; 1975.

22. Nowak MA, Tarnita CE, Wilson EO. The evolution of eusociality. Nature. 2010;466(7310):1057–62.

23. Quiñones AE, Pen I. A unified model of Hymenopteran preadaptations that trigger the evolutionary transition to eusociality. Nat Commun. 2017;8:15920.

24. Danforth BN, Conway L, Ji S. Phylogeny of eusocial Lasioglossum reveals multiple losses of eusociality within a primitively eusocial clade of bees (Hymenoptera: Halictidae). Syst. Biol. 2003;52(1):23–36.

25. Gibbs J, Brady SG, Kanda K, Danforth BN. Phylogeny of halictine bees supports a shared origin of eusociality for Halictus and Lasioglossum (Apoidea: Anthophila: Halictidae). Mol. Phylogenetics Evol. 2012;65(3):926–39.

26. Michener CD. Comparative social behavior of bees. Annu. Rev. Entomol. 1969;14(1):299–342.

27. Oster GF, Wilson EO. Caste and ecology in the social insects. Monogr Popul Biol. 1978;12:1–352.

28. Santos PKF, Arias MC, Kapheim KM. Loss of developmental diapause as prerequisite for social evolution in bees. Biology Letters 15(8):20190398.

29. Smith AR, Wcislo WT, O’Donnell S. Survival and productivity benefits to social nesting in the sweat bee Megalopta genalis (Hymenoptera: Halictidae). Behav. Ecol. Sociobiol. 2007;61(7):1111–20.

30. Yagi N, Hasegawa E. A halictid bee with sympatric solitary and eusocial nests offers evidence for Hamilton’s rule. Nat Commun. 2012;3(1):939.

31. Soucy SL. nesting biology and socially polymorphic behavior of the sweat bee Halictus rubicundus (Hymenoptera: Halictidae). Ann. Entomol. Soc. Am. 2002;95(1):57–65.

32. Davison P, Field J. Social polymorphism in the sweat bee Lasioglossum (Evylaeus) calceatum. Insect Soc. 2016;63(2):327–38.

33. Plateaux-Quénu C. Comparative biological data in two closely related eusocial species: Evylaeus calceatus (Scop.) and Evylaeus albipes (F.) (Hym., Halictinae). Insect Soc. 1992;39:351–64.

34. Yanega D. Environmental influences on male production and social structure in Halictus rubicundus (Hymenoptera: Halictidae). Insect Soc. 1993;40(2):169–80.

35. Gruber J, Field J. Male survivorship and the evolution of eusociality in partially bivoltine sweat bees. PloS One. 2022;17(10):e0276428.

36. Fisher RA. The genetical theory of natural selection. Oxford, England: Clarendon Press; 1930.

37. Trivers RL, Hare H. Haploidploidy and the evolution of the social insect. Science. 1976;191(4224):249–63.

38. Remnant EJ, Koetz A, Tan K, Hinson E, Beekman M, Oldroyd BP. Reproductive interference between honeybee species in artificial sympatry. Mol Ecol. 2014;23(5):1096–107.

39. Tsuchida K, Yamaguchi A, Kanbe Y, Goka K. Reproductive interference in an introduced bumblebee: polyandry may mitigate negative reproductive impact. Insects. 2019;10(2).

40. Gibbs T, Levin SA, Levine JM. Coexistence in diverse communities with higher-order interactions. Proc. Natl. Acad. Sci. U.S.A. 2022;119(43):e2205063119.

41. Schwarz MP, Richards MH, Danforth BN. Changing paradigms in insect social evolution: insights from halictine and allodapine bees. Annu Rev Entomol. 2007;52:127–50.

42. Field J. Patterns of provisioning and iteroparity in a solitary halictine bee,Lasioglossum (Evylaeus) fratellum (Perez), with notes on L. (E.) calceatum (Scop.) and L. (E.) villosulum (K.). Insect Soc. 1996;43(2):167–82.

43. Hennessy G, Harris C, Eaton C, Wright P, Jackson E, Goulson D, et al. Gone with the wind: effects of wind on honey bee visit rate and foraging behaviour. Animal Behaviour. 2020;161:23–31.

44. Parker D, Legg T, Folland C. A new daily central England temperature series, 1772–1991. Int. J. Climatol.1992;12:317–42.

45. Gadagkar R. Evolution of Eusociality: The advantage of assured fitness returns. Philos. Trans. R. Soc. 1990;329(1252):17–25.

46. Smith AR, Wcislo WT, O’Donnell S. Assured fitness returns favor sociality in a mass-provisioning sweat bee, Megalopta genalis (Hymenoptera: Halictidae). Behav. Ecol. Sociobiol. 2003;54:14–21.

47. Nowak MA. Five rules for the evolution of cooperation. Science. 2006;314(5805):1560–3.

48. Brede M. Short versus long term benefits and the evolution of cooperation in the prisoner’s dilemma game. PloS One.2013;8(2):e56016.

49. Fu F, Kocher SD, Nowak MA. The risk-return trade-off between solitary and eusocial reproduction. Ecol Lett. 2015;18(1):74–84.

50. Cronin A, Hirata M. Social polymorphism in the sweat bee Lasioglossum (Evylaeus) baleicum (Hymenoptera; Halictidae) in Hokkaido, Northern Japan. Insect Soc. 2003;50:379–86.

51. Thompson MJ, Jiggins CD. Supergenes and their role in evolution. Heredity. 2014;113(1):1–8.

52. Schwander T, Libbrecht R, Keller L. Supergenes and complex phenotypes. Curr Biol. 2014;24(7):R288–94.

53. Huang YC, Dang VD, Chang NC, Wang J. Multiple large inversions and breakpoint rewiring of gene expression in the evolution of the fire ant social supergene. Proc Biol Sci. 2018;285(1878).

54. Lagunas-Robles G, Purcell J, Brelsford A. Linked supergenes underlie split sex ratio and social organization in an ant. Proc. Natl. Acad. Sci. U.S.A. 2021;118(46):e2101427118.

55. Moss J.B., While G.M. 2021 The thermal environment as a moderator of social evolution. Biological Reviews 96(6), 2890–2910.

56. Ruttenberg et al., 17 Jun 2024 – preprint copy - BioRxiv

57. Purcell J, Avilés L. Gradients of precipitation and ant abundance may contribute to the altitudinal range limit of subsocial spiders: insights from a transplant experiment. Philos. Trans. R. Soc. 2008;275(1651):2617–25.

58. Soro A, Field J, Bridge C, Cardinal SC, Paxton RJ. Genetic differentiation across the social transition in a socially polymorphic sweat bee, Halictus rubicundus. Mol. Ecol. 2010;19(16):3351–63.

59. Packer L. Solitary and eusocial nests in a population of Augochlorella striata (Provancher) (Hymenoptera; Halictidae) at the northern edge of its range. Behav. Ecol. Sociobiol. 2004;27:339–44.

60. Yanega D. Social plasticity and early-diapausing females in a primitively social bee. Proc Natl Acad Sci U S A. 1988;85(12):4374–7.

61. Kocher SD, Paxton RJ. Comparative methods offer powerful insights into social evolution in bees. Apidologie. 2014;45(3):289–305.

62. Kapheim KM, Pan H, Li C, Blatti C, 3rd, Harpur BA, Ioannidis P, et al. Draft genome assembly and population genetics of an agricultural pollinator, the solitary alkali bee (Halictidae: Nomia melanderi). G3. 2019;9(3):625–34.

63. Majewski P., Lampa P., Burduk R., Reiner J. 2023 Prediction of the remaining time of the foraging activity of honey bees using spatiotemporal correction and periodic model re-fitting. Computers and Electronics in Agriculture 205:107596

64. Halsch CA, Shapiro AM, Fordyce JA, Nice CC, Thorne JH, Waetjen DP, et al. Insects and recent climate change. Proc Natl Acad Sci U S A. 2021;118(2):e2002543117.

65. Batchelder P, Kinney RO, Demlow L, Lynch CB. Effects of temperature and social interactions on huddling behavior in Mus musculus. Physiol. Behav. 1983;31(1):97–102.

66. Bartolini T, Butail S, Porfiri M. Temperature influences sociality and activity of freshwater fish. Environ. Biol. Fishes. 2015;98(3):825–32.

67. Kuruvilla M, Dell A, Olson AR, Knouft J, Grady JM, Forbes J, et al. The effect of temperature on fish swimming and schooling is context dependent. Oikos. 2023;2023(2):e09202.

